# Analysis of Shared Heritability in Common Disorders of the Brain

**DOI:** 10.1101/048991

**Authors:** V Anttila, B Bulik-Sullivan, H Finucane, R Walters, J Bras, L Duncan, V Escott-Price, G Falcone, P Gormley, R Malik, N Patsopoulos, S Ripke, Z Wei, D Yu, PH Lee, P Turley, IGAP consortium, IHGC consortium, ILAE Consortium on Complex Epilepsies, IMSGC consortium, IPDGC consortium, METASTROKE and Intracerebral Hemorrhage Studies of the International Stroke Genetics Consortium, Attention-Deficit Hyperactivity Disorder Working Group of the Psychiatric Genomics Consortium, Autism Spectrum Disorders Working Group of The Psychiatric Genomics Consortium, Bipolar Disorders Working Group of the Psychiatric Genomics Consortium, Eating Disorders Working Group of the Psychiatric Genomics Consortium, Major Depressive Disorder Working Group of the Psychiatric Genomics Consortium, Tourette Syndrome and Obsessive Compulsive Disorder Working Group of the Psychiatric Genomics Consortium, Schizophrenia Working Group of the Psychiatric Genomics Consortium, G Breen, C Churchhouse, C Bulik, M Daly, M Dichgans, SV Faraone, R Guerreiro, P Holmans, K Kendler, B Koeleman, CA Mathews, AL Price, JM Scharf, P Sklar, J Williams, N Wood, C Cotsapas, A Palotie, JW Smoller, P Sullivan, J Rosand, A Corvin, BM Neale, on behalf of the Brainstorm consortium

**Author notes:** **Author Information** Correspondence and requests for materials should be addressed to V.A., A.C. or B.M.N. One Sentence Summary: Comprehensive heritability analysis of brain phenotypes demonstrates a 59 clear role for common genetic variation across neurological and psychiatric disorders and 60 behavioral-cognitive traits, with substantial overlaps in genetic risk.

## Abstract

Disorders of the brain exhibit considerable epidemiological comorbidity and frequently share symptoms, provoking debate about the extent of their etiologic overlap. We quantified the genetic sharing of 25 brain disorders based on summary statistics from genome-wide association studies of 215,683 patients and 657,164 controls, and their relationship to 17 phenotypes from 1,191,588 individuals. Psychiatric disorders show substantial sharing of common variant risk, while neurological disorders appear more distinct from one another. We observe limited evidence of sharing between neurological and psychiatric disorders, but do identify robust sharing between disorders and several cognitive measures, as well as disorders and personality types. We also performed extensive simulations to explore how power, diagnostic misclassification and phenotypic heterogeneity affect genetic correlations. These results highlight the importance of common genetic variation as a source of risk for brain disorders and the value of heritability-based methods in understanding their etiology.

---

The classification of brain disorders has evolved over the past century, reflecting the medical and scientific communities’ best assessments of the presumed root causes of clinical phenomena such as behavioral change, loss of motor function, spontaneous movements or alterations of consciousness. A division between neurology and psychiatry developed, with the more directly observable phenomena (such as the presence of emboli, protein tangles, or unusual electrical activity patterns) generally defining the neurological disorders(1). Applying modern methods to understand the genetic underpinnings and categorical distinctions between brain disorders may be helpful in informing next steps in the search for the biological pathways underlying their pathophysiology(2, 3).

In general, brain disorders (here excepting those caused by trauma, infection or cancer) show substantial heritability from twin and family studies (4). Epidemiological and twin studies have explored patterns of phenotypic overlaps(5–7), and substantial comorbidity has been reported for many pairs of disorders, including bipolar disorder-migraine(8), stroke-major depressive disorder(MDD)(9), epilepsy-autism spectrum disorders (ASD) and epilepsy-attention deficit hyperactivity disorder (ADHD)(10, 11). Furthermore, neurological and psychiatric research has shown that mutations in the same ion channel genes confer pleiotropic risk for multiple distinct brain phenotypes(12–14). Recently, genome-wide association studies (GWAS) have demonstrated that individual common risk variants show overlap across traditional diagnostic boundaries (15, 16), and that disorders like schizophrenia, MDD and bipolar disorder can have strong genetic correlations(17).

GWAS have also demonstrated that common genetic variation substantially contributes to the heritability of brain disorders. In most cases, this occurs via many common variants, each of small effect, with examples in Alzheimer’s disease(18), bipolar disorder(19), migraine(20), Parkinson’s disease(21), and schizophrenia(22). In addition to locus discovery, the degree of distinctiveness (23) across a wide set of neurological and psychiatric phenotypes can now be evaluated with the introduction of novel heritability-based methods(24) and sufficiently large sample sizes. These analyses can shed light on the nature of these diagnostic boundaries and explore the extent of shared common variant genetic influences.

## Study design

We formed the Brainstorm consortium, a collaboration among GWAS meta-analysis consortia of 25 disorders (see Data sources), to perform the first comprehensive heritability and correlation analysis of brain disorders. We included all common brain disorders for which we could identify a GWAS meta-analysis consortium of sufficient size for heritability analysis that was willing to participate. The total study sample consists of 215,683 cases of different brain disorders and 657,164 controls (Table 1), and provides coverage of a majority of ICD-10 blocks covering mental and behavioral disorders and diseases of the central nervous system. Also included are 1,191,588 samples for 13 “behavioral-cognitive” phenotypes (n=744,486) chosen for being traditionally viewed as brain-related, and four “additional” phenotypes (n=447,102) selected to represent known, well-delineated etiological processes (e.g. immune disorders [Crohn’s disease] and vascular disease [coronary artery disease]; Table 2) or anthropomorphic measures (height and BMI).

**Table 1.** Brain disorder phenotypes used in the Brainstorm project. Indented phenotypes are part of a larger whole, e.g. the epilepsy study consists of the joint analysis of focal epilepsy and generalized epilepsy. Numbers in gray denote a sample set which is non-unique, e.g. cardioembolic stroke samples are a subset of ischemic stroke samples. ADHD – attention deficit hyperactivity disorder; OCD – obsessive-compulsive disorder. ‘Anxiety disorders’ refers to a meta-analysis of five subtypes (generalized anxiety disorder, panic disorder, social phobia, agoraphobia and specific phobias). Source details are listed under Data Sources and the references in Table S1.

| Psychiatric disorders |  |  |  | Neurological disorders |  |  |  |
| --- | --- | --- | --- | --- | --- | --- | --- |
| Disorder | Source | Cases | Controls | Disorder | Source | Cases | Controls |
| ADHD | PGC-ADD2 | 12,645 | 84,435 | Alzheimer's disease | IGAP | 17,008 | 37,154 |
| Anorexia nervosa | PGC-ED | 3,495 | 11,105 | Epilepsy | ILAE | 7,779 | 20,439 |
| Anxiety disorders | ANGST | 5,761 | 11,765 | Focal epilepsy | " | 4,601 | 17,985 |
| Autism spectrum disorder | PGC-AUT | 6,197 | 7,377 | Generalized epilepsy | " | 2,525 | 16,244 |
| Bipolar disorder | PGC-BIP2 | 20,352 | 31,358 | Intracerebral hemorrhage | ISGC | 1,545 | 1,481 |
| Major depressive disorder | PGC-MDD2 | 16,823 | 25,632 | Ischemic stroke | METASTROKE | 10,307 | 19,326 |
| OCD | PGC-OCDS | 2,936 | 7,279 | Cardioembolic stroke | " | 1,859 | 17,708 |
| PTSD | PGC-PTSD | 2,424 | 7,113 | Early-onset stroke | " | 3,274 | 11,012 |
| Schizophrenia | PGC-SCZ2 | 33,640 | 43,456 | Large-vessel disease | " | 1,817 | 17,708 |
| Tourette Syndrome | PGC-OCDS | 4,220 | 8,994 | Small-vessel disease | " | 1,349 | 17,708 |
|  |  |  |  | Migraine | IHGC | 59,673 | 316,078 |
|  |  |  |  | Migraine with aura | " | 6,332 | 142,817 |
|  |  |  |  | Migraine without aura | " | 8,348 | 136,758 |
|  |  |  |  | Multiple sclerosis | IMSGC | 5,545 | 12,153 |
|  |  |  |  | Parkinson's disease | IPDGC | 5,333 | 12,019 |
| *Total psychiatric* |  | *108,493* | *238,514* | *Total neurologic* |  | *107,190* | *418,650* |

**Table 2.**
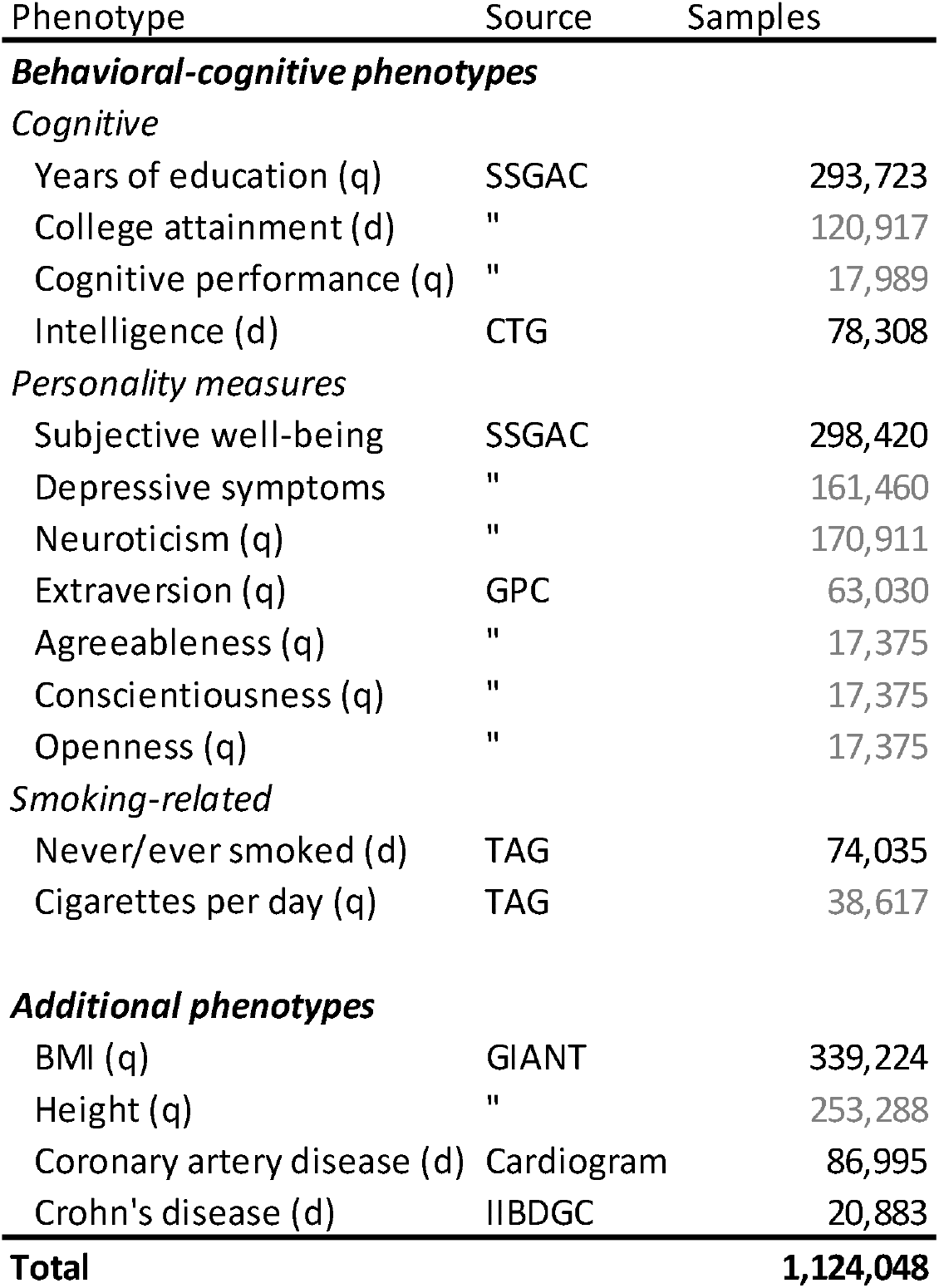
Behavioral-cognitive and additional phenotypes used in the study. Numbers in gray denote overlapping study sets, e.g. samples in the college attainment analysis are a subset of those in the analysis for years of education. (d) – dichotomous phenotype, (q) – quantitative phenotype. BMI – body-mass index. Source details are listed under Data Sources, while references are listed in Table S2.

GWAS summary statistics for the 42 disorders and phenotypes were centralized and underwent uniform quality control and processing(25). Where necessary, we generated European-only meta-analyses for each disorder to avoid potential biases arising from ancestry differences, as many of the brain disorder datasets included sample sets from diverse ancestries. Clinically relevant subtypes from three disorders (epilepsy, migraine and ischemic stroke) were also included; in these cases, the analyzed datasets are subsets of the top-level dataset, as shown in Table 1.

We have recently developed a novel heritability estimation method, linkage disequilibrium score regression (LDSC)(24), which was used to calculate heritability estimates and correlations, as well as to estimate their statistical significance from block jack-knife-based standard errors. Heritability for binary disorders and phenotypes was transformed to the liability-scale. We further performed a weighted-least squares regression analysis to evaluate whether differences relating to study makeup (such as sample size) were correlated with the magnitude of the correlation estimates. We also performed a heritability partitioning analysis using stratified LD score regression to examine whether the observed heritability was enriched in any tissue-specific regulatory partitions of the genome, using the ten top-level tissue-type and 53 functional partitions from Finucane et al. (26). Finally, simulated phenotype data was generated under several different scenarios by permuting the 120,267 genotyped individuals from the UK Biobank (25) to both evaluate power and aid in interpreting the results (see Supplementary Text).

## Heritability and correlations among brain disorders

We observed a similar range of heritability estimates among the disorders and the behavioral-cognitive phenotypes (Fig. S1A-B and Table S1, S2), roughly in line with previously reported estimates obtained from smaller datasets (see Table S3 and Supplementary Text). Three ischemic stroke subtypes (cardioembolic, large-vessel disease and small-vessel disease) as well as the “agreeableness” personality measure from NEO Five-Factor Inventory(27) had insufficient evidence of additive heritability for robust analysis and thus were excluded from further analysis(25). We did not observe a correlation between heritability estimates and factors relating to study makeup (Table S4; Fig. S1C-F). Since some of the results interpretation depends on lack of observed correlation, we explored the behavior of observed correlation vs power (Fig. S2A), standard errors (Fig. S2B) and the individual results (Fig. S2C and D) to identify where we can be reasonably robust in claiming lack of correlation with current datasets.

In expanding on the number of pairwise comparisons in brain disorders, we observed widespread sharing across psychiatric disorders (Fig. 1 and S3) beyond those previously reported (17), but not among neurological disorders. Among the psychiatric disorders, schizophrenia showed significant genetic correlation with most of the psychiatric disorders, while MDD was positively (though not necessarily significantly) correlated with every other disorder tested. Further, schizophrenia, bipolar disorder, anxiety disorders, MDD and ADHD each showed a high degree of correlation to the four others (average *r_g_* = 0.40; Table S5). Anorexia nervosa, obsessive-compulsive disorder (OCD) and schizophrenia also demonstrated significant sharing amongst themselves. On the other hand, the common variant risk of both ASD and Tourette Syndrome (TS) appear to be somewhat distinct from other psychiatric disorders, although with significant correlation between TS, OCD and MDD, as well as between ASD and schizophrenia. Post-traumatic stress disorder (PTSD) alone showed no significant correlation with any of the other psychiatric phenotypes (though some correlation to ADHD and MDD was observed, Fig. 1). The modest power of the ASD, PTSD and TS meta-analyses, however, limits the strength of this conclusion (Fig. S2C).

**Figure 1.**
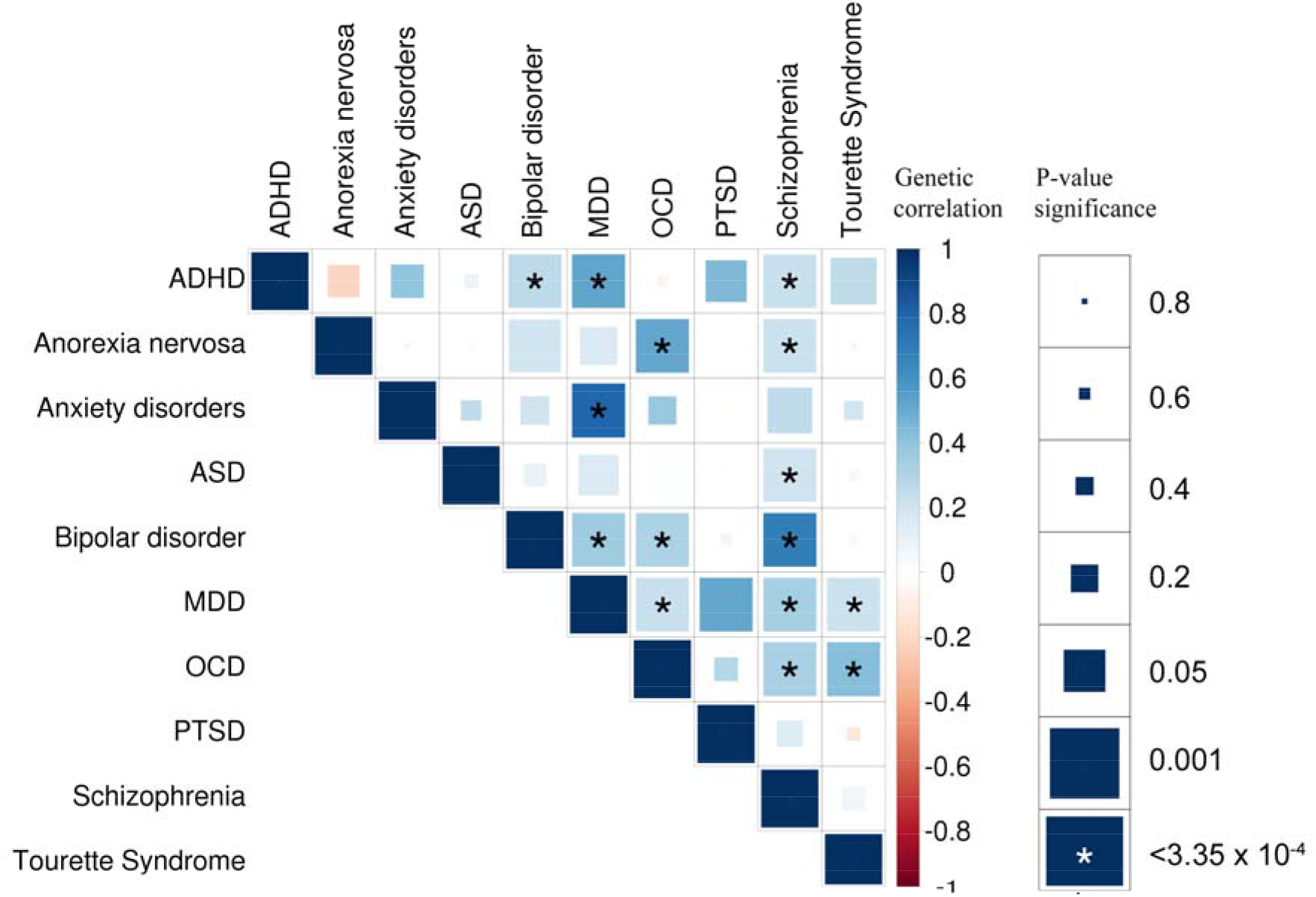
Genetic correlation matrix across psychiatric phenotypes. Color of each box indicates the magnitude of the correlation, while size of the boxes indicates its significance, with significant correlations filling each box completely. Asterisks indicate genetic correlations which are significant after Bonferroni correction. ADHD – attention deficit hyperactivity disorder; ASD – autism spectrum disorder; MDD – major depressive disorder; OCD – obsessive-compulsive disorder; PTSD – post-traumatic stress disorder.

Neurological disorders revealed greater specificity, and a more limited extent of genetic correlation than the psychiatric disorders (Fig. 2 and S4, Table S5). Parkinson’s disease, Alzheimer’s disease, generalized epilepsy and multiple sclerosis showed little to no correlation with any other brain disorders. Focal epilepsy showed the highest degree of genetic correlation among the neurological disorders (average *r_g_* = 0.46, excluding other epilepsy datasets), though none were significant, reflecting the relatively modest power of the current focal epilepsy meta-analysis (Fig. S2C). However, the modest heritability and the broad pattern of sharing observed for focal epilepsy may be consistent with considerable heterogeneity and potentially even diagnostic misclassification across a range of neurological conditions.

**Figure 2.**
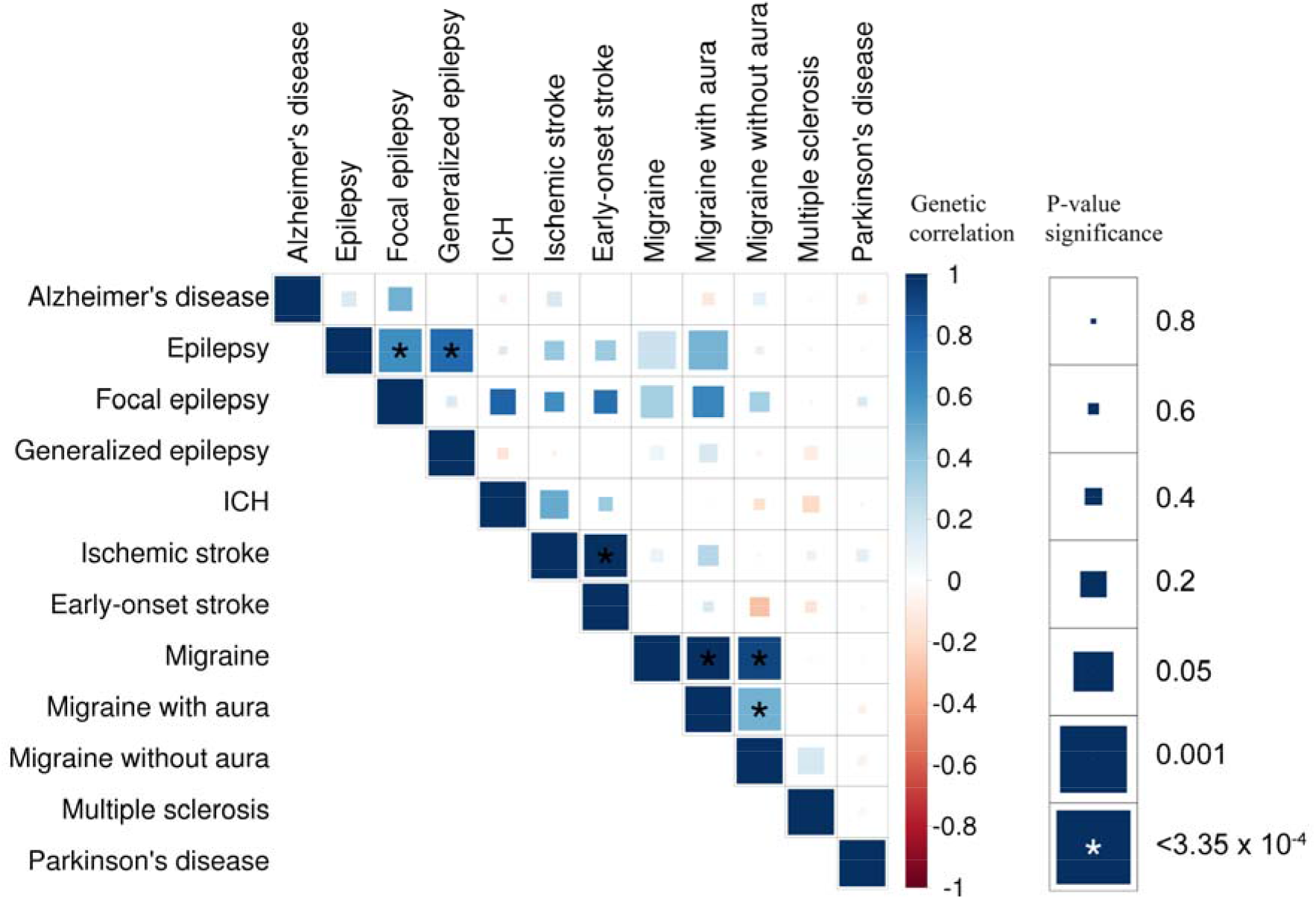
Genetic correlation matrix across neurological phenotypes. Color of each box indicates the magnitude of the correlation, while size of the boxes indicates its significance, with significant correlations filling each box completely. Asterisks indicate genetic correlations which are significant after Bonferroni correction. Some phenotypes have substantial overlaps (see Table 1), e.g. all cases of generalized epilepsy are also cases of epilepsy. Asterisks indicate significant genetic correlation after multiple testing correction. ICH – intracerebral hemorrhage.

In the cross-category correlation analysis, the overall pattern is consistent with limited sharing across the included neurological and psychiatric disorders (Fig. 3; average *r_g_* = 0.03). The only significant cross-category correlations were with migraine, suggesting it may share some of its genetic architecture with psychiatric disorders; migraine-ADHD (*r_g_* = 0.26, p=8.81 × 10^−8^), migraine-TS (*r_g_* = 0.19, p=1.80 × 10^−5^), and migraine-MDD (*r_g_* = 0.32, p=1.42 × 10^−22^ for all migraine, *r_g_* = 0.23, p=5.23 × 10^−5^ for migraine without aura, *r_g_* = 0.28, p=1.00 × 10^−4^ for migraine with aura).

**Figure 3.**
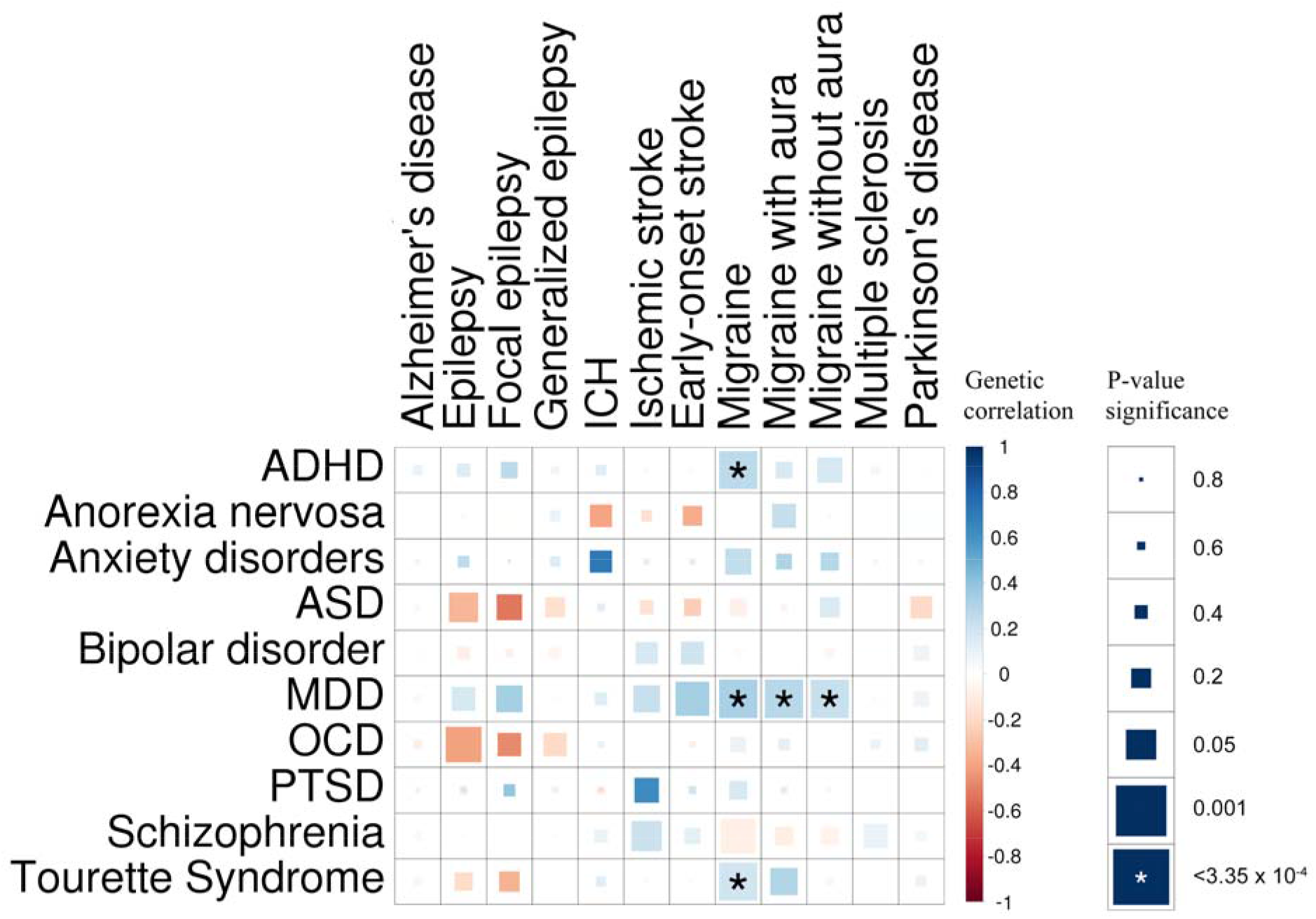
Genetic correlation matrix across neurological and psychiatric phenotypes. Color of each box indicates the magnitude of the correlation, while size of the boxes indicates its significance, with significant correlations filling each box completely. Asterisks indicate genetic correlations which are significant after Bonferroni correction. ADHD – attention deficit hyperactivity disorder; ASD – autism spectrum disorder; ICH – intracerebral hemorrhage; MDD – major depressive disorder; OCD – obsessive-compulsive disorder; PTSD – post-traumatic stress disorder.

We observed several significant genetic correlations between the behavioral-cognitive or additional phenotypes and brain disorders (Fig. 4, Table S6). Results for cognitive traits were dichotomous among psychiatric phenotypes (Fig. S5A), with ADHD, anxiety disorders, MDD and Tourette Syndrome showing negative correlations to the cognitive measures, while anorexia nervosa, ASD, bipolar disorder and OCD showed positive correlations. Schizophrenia showed more mixed results, with significantly negative correlation to intelligence but positive correlation to years of education. Among neurological phenotypes (Fig. S5B), the correlations were all either negative or null, with Alzheimer’s disease, epilepsy, ICH, ischemic stroke, early-onset stroke and migraine showing significantly negative correlations. Correlations with bipolar disorder(24), Alzheimer’s disease and schizophrenia have been previously reported(28)).

**Figure 4.**
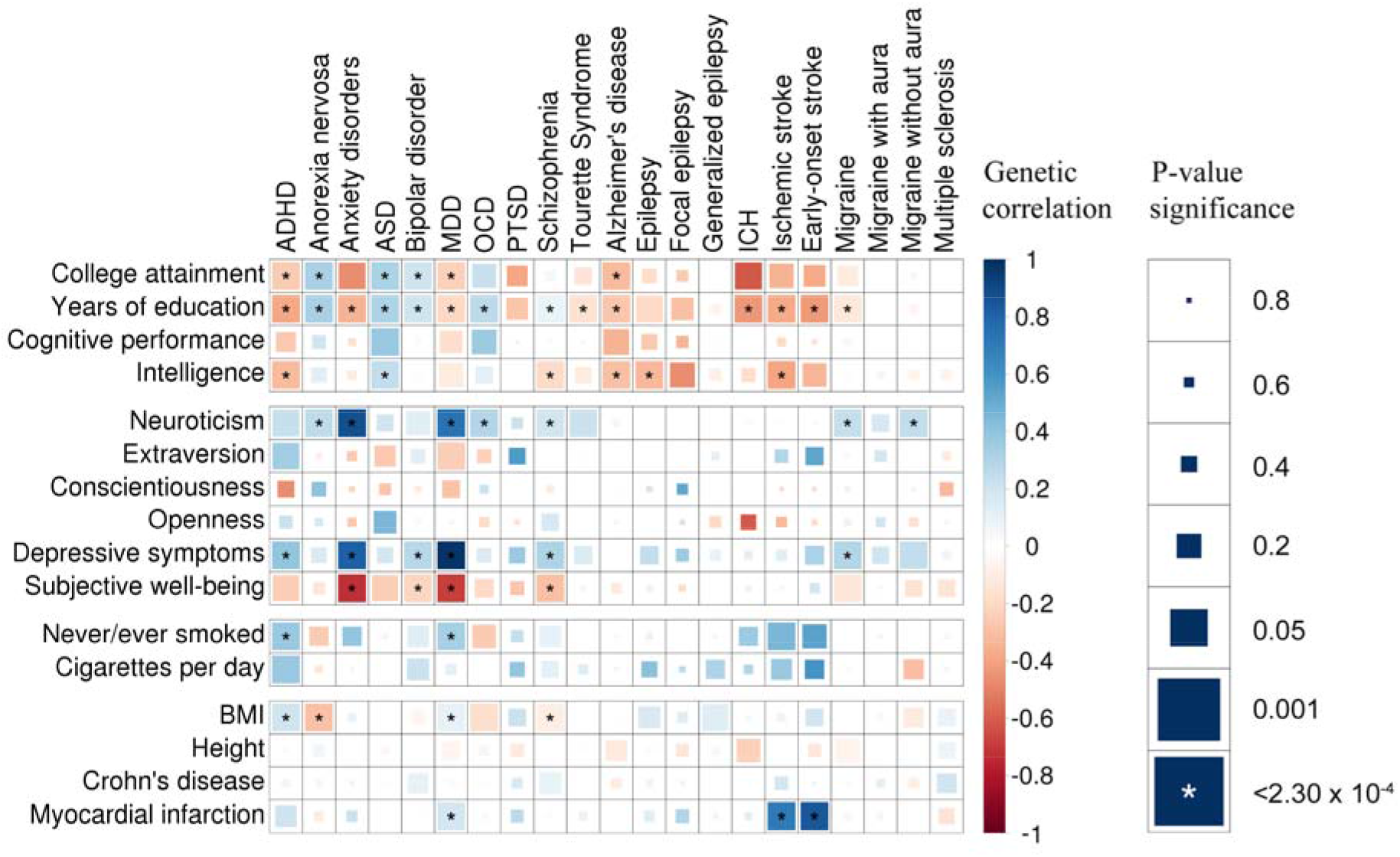
Genetic correlation matrix across brain disorders and behavioral-cognitive phenotypes. Color of each box indicates the magnitude of the correlation, while size of the boxes indicates its significance, with significant correlations filling each box completely. Asterisks indicate genetic correlations which are significant after Bonferroni correction. ADHD – attention deficit hyperactivity disorder; ASD – autism spectrum disorder; ICH – intracerebral hemorrhage; MDD – major depressive disorder; OCD – obsessive-compulsive disorder; PTSD – post-traumatic stress disorder; BMI – body-mass index.

Among the personality measures, significant positive correlations were observed for neuroticism (anorexia nervosa, anxiety disorders, migraine, migraine without aura, MDD, OCD, schizophrenia and Tourette Syndrome; Fig. S6A), depressive symptoms (ADHD, anxiety disorder, bipolar disorder, MDD, and schizophrenia) and subjective well-being (anxiety disorder, bipolar disorder, MDD, as well as replicating the previously reported correlation between neuroticism with both MDD and schizophrenia(29)). For smoking-related measures, the only significant genetic correlations were to never/ever smoked (MDD: *r_g_* = 0.33, p=3.10 × 10^−11^ and ADHD: *r_g_* = 0.37, p=3.15 × 10^−6^).

Among the additional phenotypes, the two diseases chosen as examples of disorders with well-defined etiologies had different results: Crohn’s disease, representing immunological pathophysiology, showed no correlation with any of the study phenotypes, while the phenotype representing vascular pathophysiology (coronary artery disease) showed significant correlation to MDD (*r_g_* = 0.19, p=8.71 × 10^−5^) as well as the two stroke-related phenotypes (*r_g_* = 0.69, p=2.47 × 10^−6^ to ischemic stroke and *r_g_* = 0.86, p=2.26 × 10^−5^ for early-onset stroke), suggesting shared genetic effects across these phenotype. Significant correlations were also observed for BMI, which was positively correlated with ADHD and MDD, and negatively correlated with anorexia nervosa (as previously reported with a different dataset(24)) and schizophrenia.

Our enrichment analysis (Fig. S7, Table S7 and S8) demonstrated novel significant heritability enrichments between central nervous system (CNS) and generalized epilepsy, MDD, TS, college attainment, intelligence, neuroticism, never/ever smoked); depressive symptoms and adrenal/pancreatic cells and tissues, as well as between immune system cells and multiple sclerosis. We also note with interest that the psychiatric disorders with large numbers of identified GWAS loci (bipolar disorder, MDD and schizophrenia) and the only cross-correlated neurological disorder with the same (migraine) all show enrichment to conserved regions, while the other neurological disorders with similar numbers of loci (MS and Alzheimer’s and Parkinson’s diseases) do not (Fig. S7A, B). Significant enrichment to conserved regions was also observed to neuroticism, intelligence and college attainment and to H3K9ac peaks for BMI. We also replicate the previously reported (CNS) enrichment for schizophrenia, bipolar disorder and years of education (here in a larger dataset compared to the original report, but with considerable sample overlap), and observe the previously reported enrichments for BMI (CNS), years of education (CNS), height (connective tissues and bone, cardiovascular system and other) and Crohn’s disease (hematopoietic cells) from the same datasets (Fig. S7C, D) (26).

## Discussion

By integrating and analyzing the current genome-wide association summary statistic data from consortia of 25 brain disorders, we find that psychiatric disorders broadly share a considerable portion of their common variant genetic risk, especially across schizophrenia, MDD, bipolar disorder, anxiety disorder and ADHD, while neurological disorders are more genetically distinct. Across categories, psychiatric and neurologic disorders share relatively little of their common genetic risk, suggesting that multiple different and largely independently regulated etiological pathways may give rise to similar clinical manifestations (e.g., psychosis, which manifests in both schizophrenia(30) and Alzheimer’s disease(31)). Except for migraine, which appears to share some genetic architecture with psychiatric disorders, the existing clinical delineation between neurology and psychiatry is recapitulated at the level of common variant risk for the studied disorders.

Given that the broad and continuous nature of psychiatric disorder spectra in particular has been clinically recognized for a long time(32–34) and that patients can, in small numbers, progress from one diagnosis to another(35), we evaluated to what extent diagnostic misclassification could explain the observed correlations. Genetic correlation could arise if, for example, substantial numbers of patients progress through multiple diagnoses over their lifetime, or if some specific diagnostic boundaries between phenotype pairs are particularly porous to misclassification; while it would be unlikely to observe large-scale misclassification of migraine as schizophrenia, for example, there may be more substantial misclassification between other pairs, consistent with the clinical controversies in classification. Previous work(36) suggests that substantial misclassification (on the order of 15-30%, depending on whether it is uni- or bidirectional) is required to introduce high levels of genetic correlation. We sought to confirm and expand upon these estimates by performing large-scale simulations and calculating the resulting correlations across a variety of scenarios (Fig. S8, S9, Table S9 and Supplementary Text). First, we established that the observed heritability of the simulated misclassified traits behaves as expected (Fig. S8A), and that the effects on observed correlation (Fig. S8B and S8C) are in line with the estimates from family data(36). We further explored the effect of misclassification on observed *r_g_* given the correlation observed in real data. Reasonably low levels of misclassification or changes to the exact level of heritability appear unlikely to induce substantial changes in the estimated genetic correlation, though a lower observed heritability caused by substantial misclassification (Fig. S8A) will decrease the power to estimate the genetic overlap, as observed in the power analysis (Fig. S10). Further, such evidence of genetic overlap is unlikely to appear in the absence of underlying genetic correlation (Table S10), as it is apparent that a very high degree of misclassification (up to 79%) would be required to produce the observed correlations in the absence of any true genetic correlation. Therefore, the observed correlations suggest true sharing of a substantial fraction of the common variant genetic architecture among psychiatric disorders as well as between behavioral-cognitive measures and brain disorders.

The high degree of genetic correlation among the psychiatric disorders adds further evidence that current clinical diagnostics do not reflect the underlying genetic etiology of these disorders, and that genetic risk factors for psychiatric disorders do not respect clinical diagnostic boundaries. This suggests an interconnected nature for their genetic etiology, in contrast to neurological disorders, and underscores the need to refine psychiatric diagnostics. This study may provide important ‘scaffolding’ to support a new research framework for investigating mental disorders, incorporating many levels of information to understand basic dimensions of brain function, such as through the National Institute of Mental Health’s RDoC initiative.

The observed positive genetic correlations are consistent with a few different scenarios. For example, *r_g_* may reflect the existence of some portion of common genetic risk factors conferring equal risks to multiple disorders where other distinct additional factors contribute to the eventual clinical presentation. The presence of significant genetic correlation may also reflect the phenotypic overlap between any two disorders; for example, the sharing between schizophrenia and ADHD might reflect underlying difficulties in executive functioning, which are well-established in both disorders(37). Similarly, the sharing between anorexia nervosa, OCD and schizophrenia may reflect a shared mechanism underlying cognitive biases that extend from overvalued ideas to delusions. Another scenario is that a heritable intermediate trait confers risk to multiple outcomes, thereby giving rise to the genetic correlation, as the genetic influences on this trait will be shared for both outcomes (e.g., obesity as a risk factor for both type 2 diabetes and coronary artery disease), or that even the majority of common genetic effects are shared between a pair of traits, but each individual effect may confer different degrees of risk and lead to different aggregate genetic risk profiles. While a combination of these is likely, it will become increasingly feasible to evaluate these overlaps at the locus level as more genome-wide significant loci are identified in the future.

The low correlations observed across neurological disorders suggest that the current classification reflects relatively specific genetic etiologies, although the limited sample size for some of these disorders and lack of inclusion of disorders conceived as “circuit-based” in the literature, such as restless legs syndrome, sleep disorders and possibly essential tremor, constrains the generalizability of this conclusion. Generally, this analysis recapitulates the current understanding of the relatively distinct primary etiology underlying these disorders; degenerative disorders (such as Alzheimer’s and Parkinson’s diseases) would not be expected *a priori* to share their polygenic risk profiles with a neuro-immunological disorder (like multiple sclerosis) or neurovascular disorder (like ischemic stroke). Similarly, we see limited evidence for the reported co-morbidity between migraine with aura and ischemic stroke(38) (*r_g_* = 0.29, p=0.099); however, the standard errors of this comparison are too high to draw strong conclusions. At the disorder subtype level, migraine with and without aura (*r_g_* = 0.48, p=1.79 × 10^−5^) shows substantial genetic correlation, while focal and generalized epilepsy (*r_g_* = 0.16, p = 0.388) show much less.

The few significant correlations across neurology and psychiatry, namely between migraine and ADHD, MDD and TS, suggest modest shared etiological overlap across the neurology/psychiatry distinction. The co-morbidity of migraine with MDD, Tourette Syndrome and ADHD has been previously reported in epidemiological studies (39–42), while in contrast, the previously reported co-morbidity between migraine and bipolar disorder seen in epidemiological studies (43) was not reflected in our estimate of genetic correlation (*r_g_* = −0.03, p=0.406).

Several phenotypes show only very low-level correlations with any of the other disorders and phenotypes studied here, despite large sample size and robust evidence for heritability, suggesting their common variant genetic risk may largely be unique. Alzheimer’s disease, Parkinson’s disease, and multiple sclerosis show extremely limited sharing with the other phenotypes and with each other. Neuroinflammation has been implicated in the pathophysiology of each of these conditions(44–46), as it has for migraine(47) and many psychiatric conditions, including schizophrenia(48), but no considerable shared heritability was observed with either of those conditions nor with Crohn’s disease, nor did we observe enrichment for immune-related tissues in the functional partitioning (Fig. S7) as we did for Crohn’s disease. While this observation does not preclude shared neuroinflammatory mechanisms in these disorders, it does suggest that on a large scale, common variant genetic influences on these inflammatory mechanisms are not shared between these disorders. Further, we only observed significant enrichment of heritability for immunological cells and tissues in multiple sclerosis, showing that inflammation-specific regulatory marks in the genome do not show overall enrichment for common variant risk for either Alzheimer’s or Parkinson’s diseases (though this does not preclude the effects of specific, non-polygenic neuroinflammatory mechanisms(49)). Among psychiatric disorders, ASD and TS showed a similar absence of correlation with other disorders, although this could reflect small sample sizes.

Analysis of the Big Five personality measures suggest that the current sample sizes for personality data are beginning to be sufficiently large for correlation testing; neuroticism, which has by far the largest sample size, shows several significant correlations. Most significant of these was to MDD (*r_g_* = 0.737, p=5.04 × 10^−96^), providing further evidence for the link between these phenotypes, reported previously with polygenic risk scores(50) and twin studies(51, 52); others included schizophrenia, anxiety disorders, migraine, migraine without aura, and OCD (Table S6). Further, the observation of strong correlation between MDD and anxiety disorders together with their remarkably strong and similar patterns of correlation between each of these disorders and the dimensional measures of depressive symptoms, subjective well-being, and neuroticism suggests that they all tag a fundamentally similar underlying etiology. The novel significant correlation between coronary artery disease and MDD supports the long-standing epidemiological observation of a link between MDD and CAD(53), while the observed correlation between ADHD and smoking initiation (*r_g_* = 0.374, p=3.15 × 10^−6^) is consistent with the epidemiological evidence of overlap(54) and findings from twin studies(55), supporting the existing hypothesis that impulsivity inherent in ADHD may drive smoking initiation and potentially dependence (though other explanations, such as reward system dysfunction would fit as well).

For the neurological disorders, five (Alzheimer’s disease, intracerebral hemorrhage, ischemic and early-onset stroke, and migraine) showed significant negative genetic correlation to the cognitive measures, while a further two (epilepsy and focal epilepsy) showed moderate negative genetic correlation (Fig. S5). For Alzheimer’s disease, poor cognitive performance in early life has been linked to increased risk for developing the disorder in later life(56), but to our knowledge no such connection has been reported for the other phenotypes. ADHD, anxiety disorders and MDD show a significant negative correlation to cognitive and education attainment measures, while the remaining five of the eight psychiatric disorders (anorexia nervosa, ASD, bipolar disorder, OCD, and schizophrenia) showed significant positive genetic correlation with one or more cognitive measures. These results strongly suggest the existence of a link between cognitive performance already in early life and the genetic risk for both psychiatric and neurological brain disorders. The basis of the genetic correlations between education, cognition and brain disorders may have a variety of root causes including indexing performance differences based on behavioral dysregulation (e.g., ADHD relating to attentional problems during cognitive tests) or may reflect ascertainment biases in certain disorders conditional on impaired cognition (e.g., individuals with lower cognitive reserve being more rapidly identified for Alzheimer’s disease).

BMI shows significant positive genetic correlation to ADHD, consistent with a meta-analysis linking ADHD to obesity(57), and negative genetic correlation with anorexia nervosa, OCD and schizophrenia. These results are consistent with the evidence for enrichment of BMI heritability in CNS tissues(26) and that many reported signals suggest neuronal involvement(58); this may also provide a partial genetic explanation for lower BMI in anorexia nervosa patients even after recovery(59). Given that no strong correlations were observed between BMI and any of the neurological phenotypes, it is possible to hypothesize that BMI’s brain-specific genetic architecture is more closely related to behavioral phenotypes. Ischemic stroke and BMI show surprisingly little genetic correlation in this analysis (*r_g_* = 0.07, p=0.26), suggesting that although BMI is a strong risk factor for stroke(60), there is little evidence for shared common genetic effects. These analyses also suggest that the reported reduced rates of cardiovascular disease in individuals with histories of anorexia nervosa (61, 62) are due to BMI-related effects; with the limited evidence of genetic correlation of anorexia nervosa with intracerebral hemorrhage, ischemic stroke, early-onset stroke and coronary artery disease, these results suggest that any lower cardiovascular mortality is more likely due to direct BMI-related effects rather than shared common genetic risk variants.

It is broadly apparent from the results presented here that the current clinical boundaries for the studied psychiatric phenotypes do not reflect distinct underlying pathogenic processes based on the genetic evidence, while in contrast, the studied neurological disorders show much greater genetic specificity. Although it is important to emphasize that while some disorders are under-represented here (e.g. personality disorders in psychiatry and circuit-based disorders [such as restless leg syndrome] in neurology), these results clearly demonstrate the limited evidence for widespread common genetic risk sharing between psychiatric and neurological disorders, while providing strong evidence for links between them and behavioral-cognitive measures. We highlight the need for some degree of restructuring of psychiatric nosology and that genetically informed analyses may provide a good basis for such activities, consistent with the historical knowledge from twin and family-based results. Further elucidation of individual disorders and their genetic overlap, especially as distinct loci map onto a subset of disorders and etiological processes, may form the basis for either defining new clinical phenotypes or support a move to a more continuous view of psychiatric phenotypes. Further study is needed to evaluate whether overlapping genetic contributions to psychiatric pathology may influence optimal treatment choices. Ultimately, such developments give hope to reducing diagnostic heterogeneity and eventually improving the diagnostics and treatment of psychiatric disorders.

## Acknowledgements

The authors wish to acknowledge Rosy Hoskins, Jaana Wessman and Joanna Martin for their cromulent comments on the manuscript, Matthew Whittall for inspiration, and the patients and participants of the respective consortia. For study-specific acknowledgments, see Supplementary Materials. GWAS summary statistics used in the paper are available either directly from, or via application submitted in, the web addresses listed below. Data on coronary artery disease has been contributed by CARDIoGRAMplusC4D investigators and have been downloaded from www.CARDIOGRAMPLUSC4D.ORG. matSpD is available at neurogenetics.qimrberghofer.edu.au/matSpD/. This research has been conducted using the UK Biobank Resource (application #18597).

## Data sources

**Disorder or phenotype – Consortium or dataset identifier – web address:**

### Psychiatric disorders

ADHD – PGC-ADD2 - http://www.med.unc.edu/pgc/results-and-downloads

Anorexia nervosa(63) – PGC-ED - http://www.med.unc.edu/pgc/results-and-downloads

Anxiety disorder(64) – ANGST - http://www.med.unc.edu/pgc/results-and-downloads

Autism spectrum disorders(65) – PGC-AUT - http://www.med.unc.edu/pgc/results-and-downloads

Bipolar disorder – PGC-BIP2 - http://www.med.unc.edu/pgc/results-and-downloads (soon)

Major depressive disorder – PGC-MDD2 - http://www.med.unc.edu/pgc/results-and-downloads (soon)

OCD – PGC-OCDTS - http://www.med.unc.edu/pgc/results-and-downloads

PTSD – PGC-PTSD - http://www.med.unc.edu/pgc/results-and-downloads

Schizophrenia(22) – PGC-SCZ2 – http://www.med.unc.edu/pgc/results-and-downloads

Tourette Syndrome – TSAIGC – http://www.med.unc.edu/pgc/results-and-downloads

### Neurological disorders

Alzheimer’s disease(18) – IGAP - http://www.pasteur-lille.fr/en/recherche/u744/igap

Epilepsy and subtypes, focal and generalized(66) – ILAE – http://www.epigad.org/page/show/gwas_index

Intracerebral hemorrhage(67) – ISGC - http://www.strokegenetics.com/

Ischemic stroke and subtypes (cardioembolic, early-onset, small-vessel and large-vessel)(68) – METASTROKE dataset of the ISGC – http://www.strokegenetics.com/

Migraine and subtypes, migraine with and without aura – IHGC – www.headachegenetics.org

Multiple sclerosis(69) – IMSGC - http://eaglep.case.edu/imsgc_web

Parkinson’s disease(21) – IPDGC – www.pdgene.org

### Behavioral-cognitive phenotypes

College attainment, years of education(70) – SSGAC – http://www.thessgac.org/data

Childhood cognitive performance(71) – SSGAC – http://www.thessgac.org/data

Extraversion, agreeableness, conscientiousness and openness (27) – GPC – http://www.tweelingenregister.org/GPC/

IQ(72) – CTG - http://ctg.cncr.nl/software/summary_statistics

Neuroticism, depressive symptoms and subjective well-being (73) – SSGAC - http://www.thessgac.org/data

Never/ever smoked, cigarettes per day(74) - TAG - http://www.med.unc.edu/pgc/results-and-downloads

### Additional phenotypes

BMI(58) – GIANT – https://www.broadinstitute.org/collaboration/giant

Height(75) – GIANT – https://www.broadinstitute.org/collaboration/giant

Crohn’s disease(76) – IIBDGC - http://www.ibdgenetics.org/downloads.html

Coronary artery disease(77) – Cardiogram – http://www.cardiogramplusc4d.org/downloads/

## Supplementary Materials

Materials and methods

Supplementary Text

Comparison with previous heritability estimates
Effect of phenotypic misclassification
Study-specific acknowledgements
Consortium memberships

Figures S1-10

Tables S1-10

